# Host-age prediction from fecal microbiome composition in laboratory mice

**DOI:** 10.1101/2020.12.04.412734

**Authors:** Adrian Low, Melissa Soh, Sou Miyake, Henning Seedorf

## Abstract

The life-long relationship between microorganisms and hosts has a profound impact on the overall health and physiology of the holobiont. Changes in microbiome composition throughout the lifespan of a host remain, however, largely understudied. In this study, the fecal microbiome of conventionally raised C57BL/6J mice was analyzed throughout almost the entire expected lifespan, from ‘maturing’ (9 weeks) until ‘very old’ age (112 weeks). Analysis of alpha and beta diversity suggests that gradual microbiome changes occur throughout the entire murine life but appear to be more pronounced in ‘maturing’ to ‘middle-aged’ phases. Phylum-level analysis indicates a shift in the Firmicutes/Bacteroidetes ratio in favor of the Firmicutes in the second year of adulthood. Varying successional patterns throughout life were observed for many Firmicutes OTUs, while relative abundances of Bacteroidetes OTUs varied primarily in the early life phases. Microbiome configurations at given time points were used as training sets in a Bayesian model, which in turn effectively enabled the prediction of host age. The fecal microbiome composition may therefore serve as an accurate biomarker for aging. This study further suggests that age-associated compositional differences may have considerable implications for the interpretation and comparability of animal model-based microbiome studies.

**Importance:** The life-long relationship between microorganisms and hosts has a profound impact on the overall physiology of the holobiont. Understanding the extent of gut microbiome compositional changes over the expected mouse lifespan may allow to better understand the interplay of microbiome and the host at the different life stages. In this study, we performed a two-year longitudinal study of murine fecal microbiome. Using fine-scale microbiome profiling we were able to predict the host age from the fecal microbiome composition. Moreover, we observed that the rate of compositional change appears to slow with age. The description of the compositional changes in commonly used C57BL/6J mice can be used to optimize selection of age-associated mouse models and highlights the use of microbiome-profiling as biomarker for aging.

## Introduction

The gut microbiome is known to exert wide-ranging effects on host health (1). As such, understanding the dynamics of the murine gut microbiome is important for mouse model-based research. Studies have shown that humans and mice harbor distinct gut microbiomes at different phases of their lives (2, 3). While compositional variability can be attributed to factors such as housing and diet (4, 5), age related factors such as host immunity-gut microbiota interactions are more likely to affect gut homeostasis and host health under controlled conditions (6). Over a mouse lifespan, other age-related changes include behavior (7), physiology (8), cellular biochemistry and susceptibility to diseases (9, 10). Notwithstanding, most researchers cite practical reasons over host biology when selecting younger mice for research (11).

The temporal changes within the gut microbiome of conventionally raised mice are neither well characterized nor understood thus far (11). Longitudinal studies are rarely performed compared to cross-sectional studies, partly due to the convenience of obtaining mice of different ages and accumulative cost of mice maintenance. Cross-sectional studies of the murine gut microbiome are generally focused on the early or later years of the murine life (2, 12, 13). One such study examined the gut microbiomes of ‘young’ (24 weeks old), ‘middle age’ (84 weeks old) and ‘very old’ (122 weeks old) female C57BL/6J mice and observed major shifts in nine of the most abundant bacterial families and functional genes with age and frailty (2). These shifts indicate that life-stage specific microbiome compositions could potentially also serve as biomarker of the host age. Nevertheless, the reproducibility of these shifts remains uncertain due to small sample size and high inter-individual variability, which may result in diverging microbiomes among mice of different batches (2, 4).

This study aims to elucidate the temporal changes in the gut microbiome of conventionally raised and widely used adult C7BL/6J mice. The longitudinal analysis of the murine gut microbiome over the course of its entire adult lifespan provides a highly resolved compositional profile, indicates life-stage specific microbiome compositions and may allow for more specific selection of mouse models for research questions relevant to the host age.

## Results

### Microbiome composition changes throughout life

The fecal microbiomes of nine-week-old C57BL/6J mice were characterized at regular intervals over 103 weeks (Figure 1a for experimental timeline and life phases and Supplementary Figure S1 for survival curve). Alpha and beta diversities between life phases, as well as individual timepoints, were compared to evaluate if general changes in diversity could be observed throughout the lifespan of the mice. The temporal changes in alpha diversity were compared using Shannon, Simpson, Chao1 and Pielou’s evenness indices (Figure 1b for life phases and Supplementary Figure 2 for fine scale). Shannon, Simpson’s diversity indices and Pielou’s evenness index showed similar trends. OTU diversity significantly increased as the mice aged from ‘maturing’(MR) to ‘mature’(MA) (FDR-corrected *P*-values < 0.01; Wilcoxon test; Supplementary Table S3), while no significant differences were observed between ‘MA’ to ‘middle age’(MD), ‘MD’ to ‘old’(OD) and ‘OD’ to ‘very old’(VO) (Figure 1b). However, the trend was different for Chao1 index, where significant differences (FDR-corrected *P*-values < 0.05; Wilcoxon test) were observed between ‘MA’ vs ‘MD’ and ‘MD’ vs ‘OD’ (Figure 1b). Collectively, the indices indicate that species evenness was the primary change between ‘MR’ to ‘MA’ and species richness was accountable for the change between ‘MA’ to ‘OD’. Comparison of incremental timepoints revealed gradual changes in Shannon and Simpson diversity and Pielou’s evenness with significant differences observed between five or fewer pairs of incremental timepoints (e.g. 38 vs 43 weeks, 47 vs 52 weeks, 52 vs 56 weeks old, 77 vs 82 weeks and 82 vs 86 weeks for Shannon diversity) (Supplementary Figure S2; Supplementary Table S3b). Longitudinal change in Chao1 index between incremental timepoint was indistinguishable between all but one pair of timepoints that showed a significant difference (104 vs 108 weeks) (Supplementary Figure S2).

**Figure 1.**
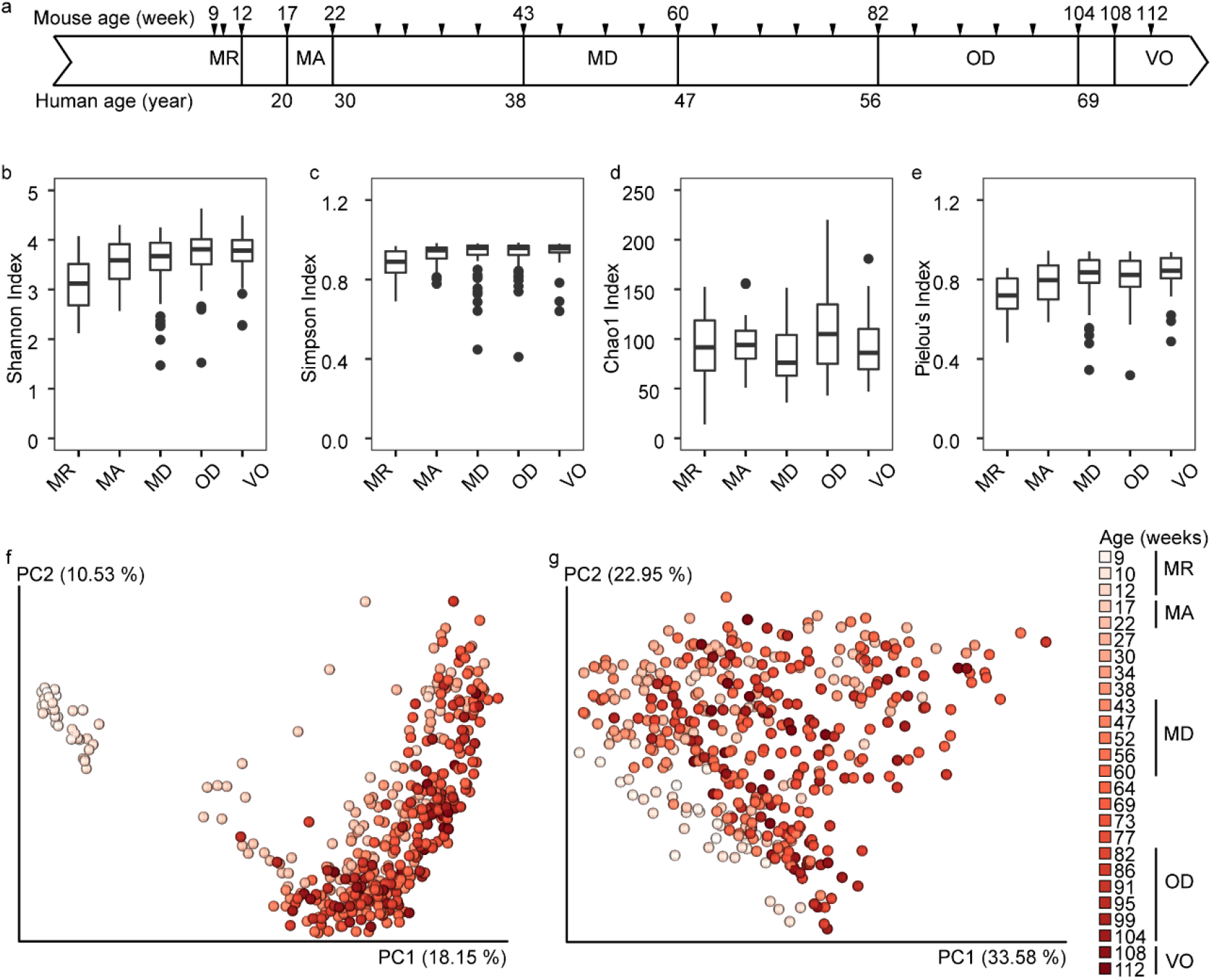
Schematic outline, changes to alpha- and beta-diversities of adult mice as they aged from 9 to 112 weeks old. Experimental timeline **(a)**. Tick marks indicate sample timepoints that began as C57BL/6J mice were 9 weeks old and ending at 112 weeks old. Life phases of mice and human equivalent age (year) based on previous reports are shown [8,10]. Boxplots of Shannon **(b)**, Simpson’s **(c)**, Chao1 **(d)** and Pielou’s **(e)** indices grouped bymouse life phases Pairwise Wilcoxon test was performed between incremental pairs of timepoints. Statistical significance is denoted with asterisks (*FDR-corrected *P* < 0.05, **FDR-corrected *P* < 0.01, ***FDR-corrected *P* < 0.001). NS is non-significant difference. ‘MR’ denotes ‘maturing’, ‘MA’ denotes ‘mature’, ‘MD’ denotes ‘middle age’, ‘OD’ denotes ‘old’ and ‘VO’ denotes ‘very old’. PCoA plots showing principal coordinates (PC) 1 and 2 of **(f)** Bray-Curtis dissimilarity and **(g)** weighted-UniFrac distance matrices. Percentage variation is shown in parenthesis. Beta-diversity measures were based on an OTU table rarefied to 3,422 counts per sample.

Consistent with this finding, there was a statistically significant difference in beta-diversity between ‘maturing’ and ‘mature’ with significant difference (FDR-corrected *P*-values < 0.001; Permutational multivariate analysis of variance (PERMANOVA)) for Bray-Curtis (Supplementary Table S3c) and weighted-UniFrac (Supplementary Table S3d) compared to later phases as the mice aged (Supplementary Figure S3). The microbiota continued to change compositionally between consecutive phases of life with statistical significance (FDR-corrected *P*-values < 0.001 for both matrices; PERMANOVA) except between ‘OD’ and ‘VO’ (Supplementary Tables S3c,d). Longitudinal comparison at closer timepoints revealed that the gut microbiotas from 56 week-old to 112 week-old mice were more stable, with decreasing number of significantly different microbiota between incremental timepoints (six pairs with FDR-corrected *P*-values < 0.05 for both matrices; PERMANOVA; Supplementary Tables S3e,f) compared to ages before 56 weeks (Figure 1c). Based on Bray-Curtis dissimilarity and weighted-UniFrac distance matrices, there was a decreasing frequency of significantly different pairs of timepoints as the mice aged (Supplementary Figures S6). Taken together, the results indicate that the murine gut microbiota underwent more compositional changes in the first year compared to the second year.

Differences in microbiome composition could be observed at different taxonomic levels. Figure 2a shows the relative abundance of the major OTUs (≥0.5% mean relative abundance) of the seven phyla over the 103-week study (*n* = 26 timepoints). Out of 1,092 OTUs, 9.3% (101 OTUs) accounted for the major OTUs. Firmicutes and Bacteroidetes constituted the predominant phyla (combined relative abundance = 96.43% ± 1.03 (mean ± SD)) over the entire time course, especially from ‘MA’ phase (17-and 22-week-old mice) to ‘MD’ (47 weeks old) where the phylum Bacteroidetes was generally more abundant (Figure 2b). Firmicutes started to comprise consistently higher relative abundances than Bacteroidetes from approximately 60 weeks (Figure 2b). Similarly, most less abundant phyla (≤ 3% mean relative abundance) were detectable at higher relative abundance in middle-aged and older mice (see Supplementary Information for details and Supplementary Figure S4).

**Figure 2.**
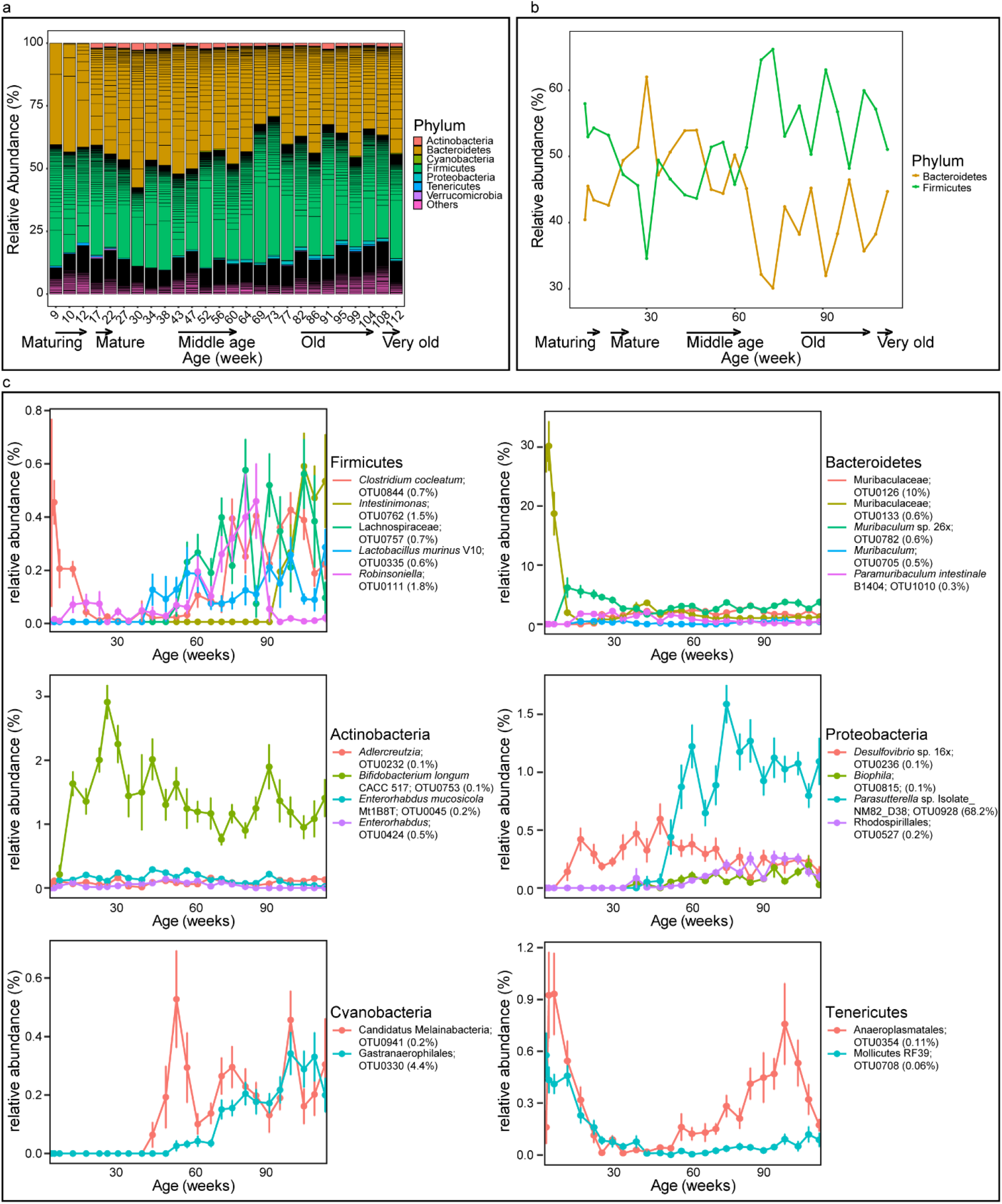
Longitudinal changes in relative abundances of bacterial taxa. **(a)** A stacked plot showing phylum groupings of major OTUs (≥0.5%). OTUs <0.5% are grouped as “others”. Each horizontal black line denotes an OTU classified at 99% similarity cut-off. **(b)** A line graph showing the change in mean relative abundance (%) of Firmicutes and Bacteroidetes over the 103 week study. **(c)** Mean relative abundances of OTUs with the top 5 importance score using a random forest regression method. Importance score in percentages are shown in parenthesis. Actinobacteria, Cyanobacteria and Tenericutes are represented by fewer than 5 predictive OTUs. Data points and error bars are the mean ± standard error of the mean, respectively. Relative abundances are based on an OTU table rarefied to 3,422 counts per sample. Taxonomic assignments >99% nucleotide identity for species and 95%-99% identity for genus level were based on top BLASTn hits and <95% nucleotide identity for family level were based on the SILVA SSU database 132 release.

OTU-level analysis revealed that only 20% (22 of the 110 OTUs) of the most abundant OTUs were detected at all analyzed timepoints (Figure 3; Supplementary Table S1). Their relative abundances differed across life stages. Four OTUs that were among the more abundant ones during the ‘MR’ stage, namely OTU0133 (closest BLAST hit (% 16S rRNA gene identity): *Paramuribaculum intestinale* B1404 (92.6%); 24.2% relative abundance), OTU0905 (*Turicibacter* sp. TS-3 (100%); 6.8% relative abundance), OTU0255 (*Lactobacillus johnsonii* G2A (99.4%); 6.6% relative abundance) and OTU0727 (*Anaerotignum* sp. 1XD42-85 (99.4%); 1.2% relative abundance), but all decreased to approximately 1% relative abundance at ‘MA’ until ‘very old (VO)’ stage (Supplementary Table S1). OTU0255 (*Lactobacillus johnsonii* G2A (99.4%)) was solely responsible for changes to *Lactobacillaceae* family over time. OTU0610 (*Faecalibaculum rodentium* Alo17 (99.4%)) stood out as the predominant OTU for the majority of adulthood with mean relative abundances of 15.5% to 17.9% between ‘MR’ to ‘VO’ phase (Supplementary Table S1). The remaining 18 OTUs observed at all timepoints stayed relatively constant throughout the mouse adult lifespan.

**Figure 3.**
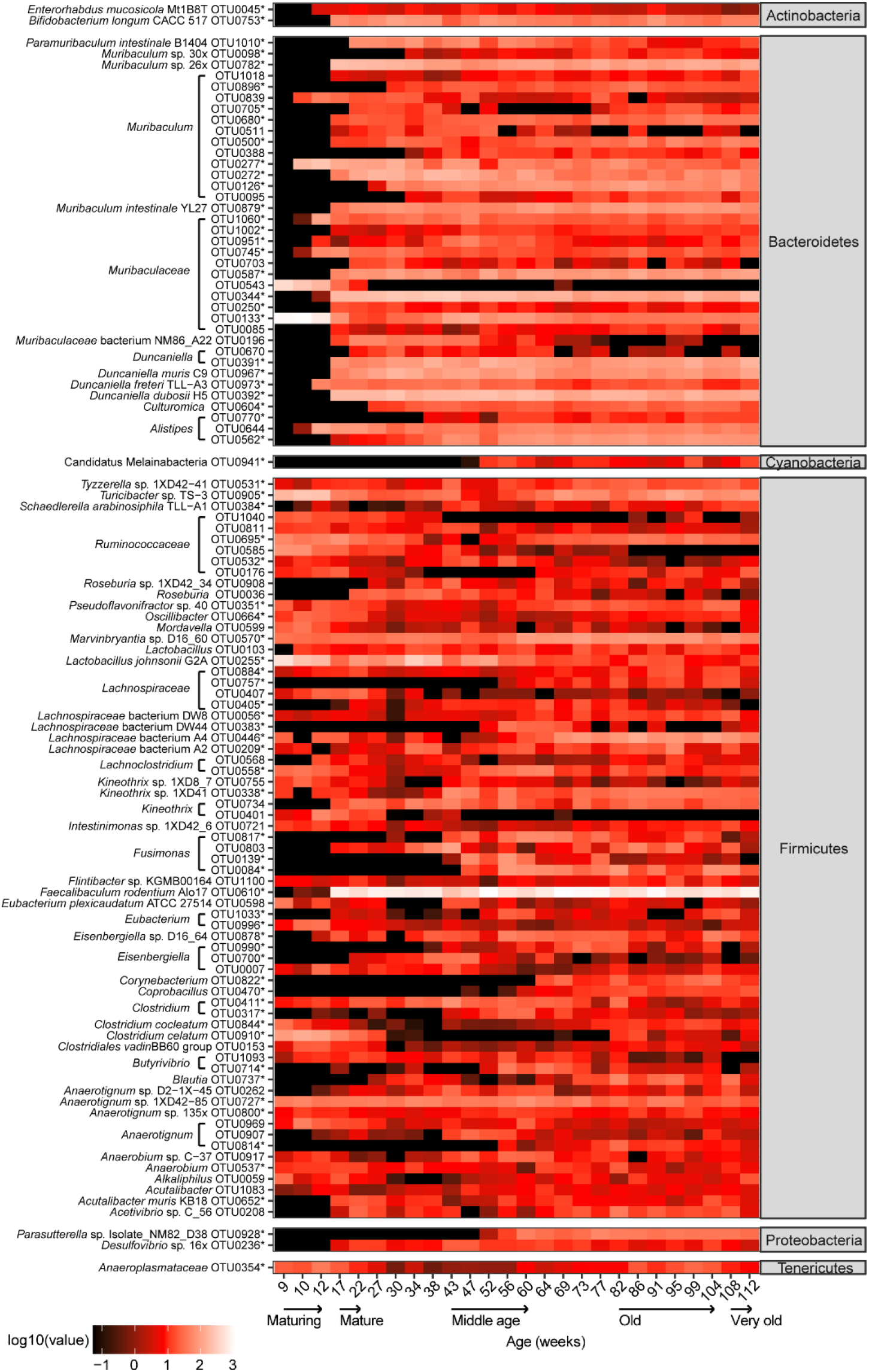
Normalized counts of 110 OTUs that have the highest mean over the 103-week-study. Counts are based on a rarefied OTU table of 3,422 counts per sample. Asterisks indicate predicted OTUs using the random forest regression method.

Successional patterns of OTUs within different phyla could be observed. The majority of Bacteroidetes OTUs were only detectable from the beginning of the ‘MA’ phase and were largely present until the mice were 112 weeks old (Figure 2a). This is in contrast with the majority of Firmicutes OTUs that were detectable at 9 weeks but have different succession patterns, e.g. a few OTUs decreased to low relative abundances as the mice aged (Figure 2a). Actinobacteria, represented by two distinct OTUs had similar succession patterns to the majority of Bacteroidetes OTUs (Figure 2a). Cyanobacteria, represented by OTU0941 (*Candidatus* Melainabacteria bacterium MEL.A1 (96.4%)), were detected at 52 weeks and remained at similar levels until 112 weeks (Figure 2a). Proteobacteria, represented by two OTUs showed dissimilar succession patterns with OTU0928 (*Parasutterella* sp. Isolate_NM82_D38 (99.4%)) detected later in the ‘MD’ phase compared to OTU0236 (*Desulfovibrio* sp. 16x (99.4%)) detected earlier at ‘MA’ phase. Tenericutes, represented by a single long-term observable bacterial OTU0354 (*Anaeroplasma abactoclasticum* 6-1 (92.1%)), had lowest mean relative abundance (0.07%) at ‘MD’ compared to other phases ranging from 0.23%-0.88% (Figure 2c; Supplementary Table S1).

### Identification of OTUs predictive for host age

The observation of the successional OTUs prompted the development of a random forest regression model to identify OTUs predictive of the 26 analyzed timepoints (14). This model showed a strong correlation (*r* = 0.958; *P* < 0.001) between predicted and actual microbiota for each timepoint. Of the 100 predictive OTUs, just five OTUs (OTU0928, OTU0126, OTU0330, OTU0111 and OTU0762) have a cumulative importance score of 86%, indicating that these OTUs have the greatest effect on the regression model (Supplementary Table S2a). Furthermore, 28 predictive OTUs of low abundance were identified (Supplementary Figure S5), of which two OTUs, OTU0232 (closest BLAST hit (% 16S rRNA gene identity): *Adlercreutzia* sp. D16-63 (95.5%)) and OTU0708 (Acholeplasmatales bacterium oral (91.8%)) were detectable at all timepoints (Supplementary Table S2b). The mean relative abundances of the top three to five OTUs of each phylum ranked by importance score over time are shown in Figure 2c. Predictive Firmicutes OTUs varied in their successional patterns: OTU0111 (*Robinsoniella sp*. D2-1X-13 (95.5%)) was more pronounced in earlier stages but diminished during ‘OD and ‘VO’ stages, while in contrast, three OTUs (OTU0844 (*Clostridium cocleatum* (100%)), OTU0757 (*Syntrophococcus* sp. BS-2 (92.6%)) and OTU0335 (*Lactobacillus murinus* V10 (99.4%)) were mid to late successors, and OTU0762 (*Intestinimonas butyriciproducens* DSM 104946 (95.9%)) was only detectable during the late phases (Figure 2c). Of the Bacteroidetes OTUs, three OTUs (OTU1010 (*Paramuribaculum intestinale* (99.4%)), OTU0705 (*Muribaculum* sp. J10 (96.0%)) and OTU0782 (*Muribaculum* sp. 26x (99.4%)) became detectable from ‘MA’ phase and remained relatively similar throughout (Figure 2c). The predictive Actinobacteria OTUs had a similar succession pattern, becoming detectable from 12 weeks old but with different relative abundances (Figure 2c). Among the Proteobacteria OTUs, OTU0236 (*Desulfovibrio* sp. 16x (99.4%)) was an early successor, OTU0928 (*Parasutterella* sp. Isolate_NM82_D38 (99.4%)) was a mid-successor, and OTU0815 (*Bilophila wadsworthia* Marseille-AA00033 (94.9%)) and OTU0527 (*Dongia* sp. CON-65 (93.4%)) were late successors (Figure 2c). The two Cyanobacteria OTUs (OTU0941 and OTU0330) have closest 16S rRNA gene identity (96.4% and 92.4%, respectively) to *Candidatus* Melainabacteria bacterium MEL.A1 were mid-successors (Figure 2c). Two Tenericutes OTUs present from the beginning decreased to low relative abundance between the ‘mature’ and ‘middle age’ phases, before OTU0354 (*Anaeroplasma abactoclasticum* 6-1 (92.1%)) increased again in relative abundance at ‘old’ phase (Figure 2c).

### Prediction of host age based on fecal microbiome composition

The observed successional pattern during the aging process and the identification of predictive OTUs for specific timepoints prompted the question: can the animal age be inferred from fecal microbiome composition? For this purpose the SourceTracker software, originally designed to use Bayesian statistics to determine the contribution of ‘source’ communities to ‘sink’ communities for monitoring water quality, was applied to our longitudinal study (15). Selection of samples (*n* = 7) as ‘source’ communities representative of the five life phases was based on timepoints with the fewest number of significantly different microbiomes as guided by pair-wise permutational multivariate analysis of variance (PERMANOVA) tests of Bray-Curtis dissimilarities (Supplementary Figure S6). The analysis revealed that specific life phases can in principle be predicted from fecal microbiomes (Figure 4a). An approximate age prediction for individual timepoints was performed by using the sum of the products of predicted life stage fractions and midpoints of life stages:

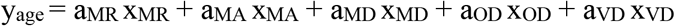

where y_age_ is the predicted age of the mouse based on fecal microbiome composition; a is the midpoint of the life stage in weeks old (i.e., ‘MR’ = 10.5, ‘MA’ = 19.5, ‘MD’ =51.5, ‘OD’ = 93 and ‘ VO’ = 110); and x is the predicted proportion of the life stage. The prediction was found to be highly accurate for younger animals (9-47 weeks old Spearman ρ = 0.925). As expected, the level of accuracy decreased for animals that are older (52-112 weeks old Spearman ρ = 0.774) (Figure 4b). This is in line with the PERMANOVA analysis that showed that the microbiome is becoming more stable in late stages of adulthood (Supplementary Figure S6).

**Figure 4.**
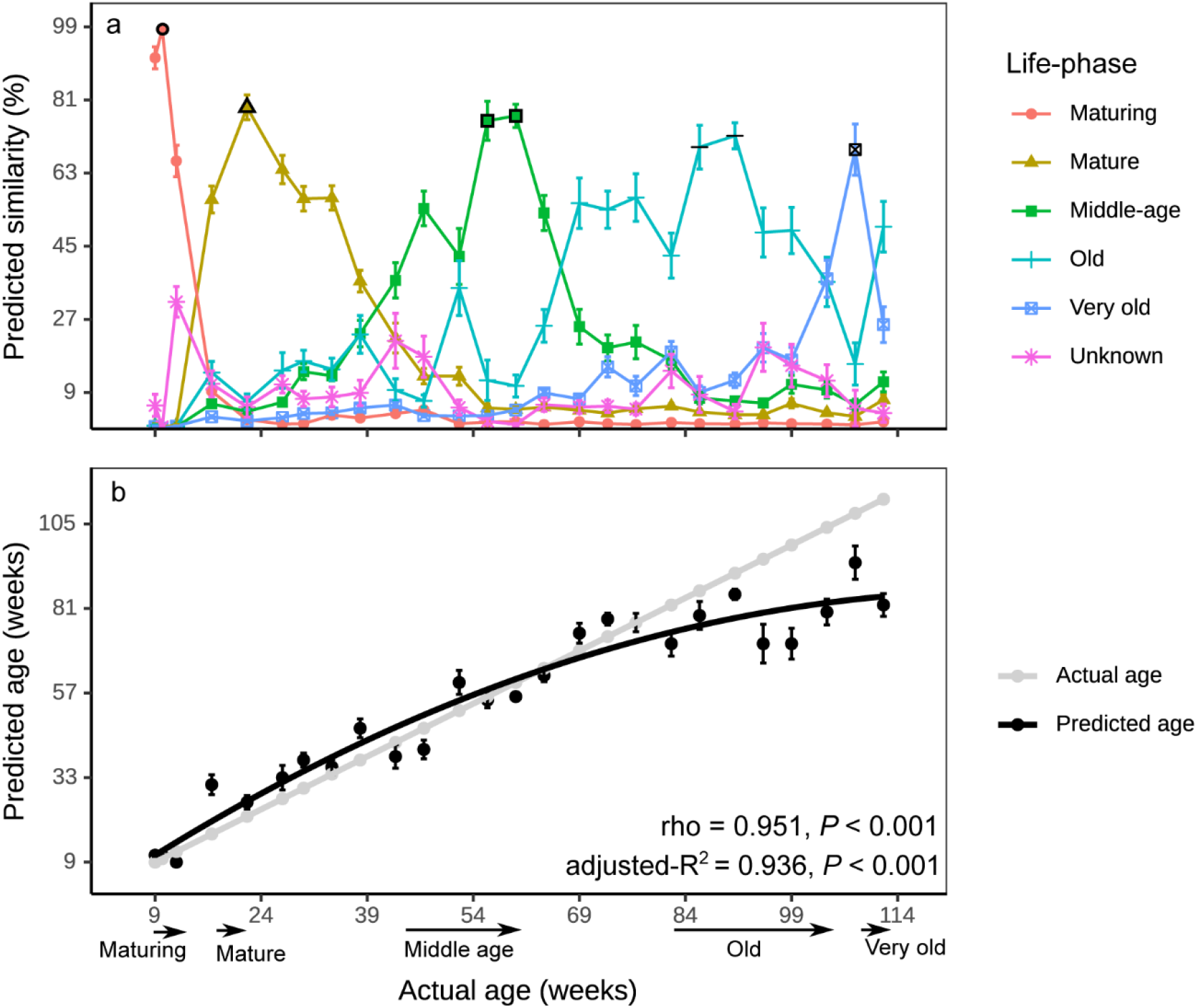
Host life-phase and age estimation based on fecal microbiota composition. **(a)** SourceTracker based prediction of probabilities for each time-point to the five life phases. Symbols in black outline show the timepoints for life phases used as ‘source’. **(b)** Correlation of predicted age to actual mouse age. The Spearman correlation coefficient (rho), adjusted-R^2^ value of the polynomial regression and thir *P*-values are shown.

## Discussion

It has been well established that the microbiomes of humans and animals change significantly in early life phases, e.g. after the weaning transition from milk to solid foods (12, 16, 17), but the longitudinal changes of the gut microbiome throughout adulthood are less well characterized. In this study, we characterized the gut microbiome of conventionally raised adult male C57BL/6J mice at regular fine-scale intervals for two years, which corresponds to the approximate lifespan of this species (10). We observed microbiome variations during murine gut development not reported in previous cross-sectional or shorter longitudinal murine studies (2, 18, 19), such as a significant increase in species richness and evenness during the early stages between ‘MR’ and ‘MA’ phase where shifts in abundance between phyla were observed (Figure 2a), but also in the later phases. Interestingly, a similar increase in alpha-diversity of swine gut microbiome was also observed during the growing years (20, 21), indicating that it may be an evolutionary trait that is conserved across species. The increased diversity also shows that the adult gut microbiome continues to change significantly after reaching host maturity between 12 to 17 weeks old. This has implications for research using mouse models. A recent meta-analysis showed that the vast majority of mice used for experiments is aged between 8-12 weeks old (11). In view of the current study, mice age should be selected carefully based on the research question.

Another notable observation is that the rate of change did not appear to be constant. There was a higher frequency of compositional change to the gut microbiome in the first year, while the second year is marked by a reduced frequency of compositional change, especially at the lower taxonomic levels (i.e. fewer OTUs in succession and more OTUs in decline) from ‘OD’ to ‘VO’ phase. At this stage, the host may experience a decreased rate of glucose and fatty acid metabolism and reduced energy expenditure (22). Feed intake (normalized to bodyweight) has been shown to not differ significantly in mice from mature to very old age (23), which indicates that the feed intake has only limited effect as underlying cause for the observed changes. This could explain the absence of significant difference in alpha-diversity between ‘OD’ and ‘VO’ mice (Figure 1). While it is generally agreed that a loss of diversity at old age may result in undesirable phenotypes such as inflammation or frailty in mice (2, 24) and increased frailty and reduced cognitive performance in humans (25, 26), our study suggests that loss of gut diversity may not necessary be associated with chronological age under controlled conditions.

Our analyses have revealed similar (e.g. Actinobacteria and Bacteroidetes) and variable (e.g. among Firmicutes OTUs) succession patterns as well as consistently observable taxa among the different phylogenetic lineages (Supplementary Tables S1 and S2b). Firmicutes remained the predominant phylum throughout ‘OD’ and ‘VO’ phases. However, prior to the late stage shift in relative abundance, Bacteroidetes relative abundance was higher than that of Firmicutes (Figure 2b). This observation in mice is in contrast with previous studies on elderly humans that generally reported an age-related decrease in the Firmicutes to Bacteroidetes ratio (27, 28). It is currently not possible to determine if these divergent patterns are specific to the host species, as has been shown for other factors such as immune maturation or microbe-host interaction (29, 30), or whether it may potentially be an artefact, e.g. resulting from cross-sectional analyses. Comparisons of our data to other murine studies may be hampered, since compositional differences may be influenced by many factors including housing facilities (4).

In our study, Firmicutes was the phylogenetically most diverse phylum, with the majority of OTUs detectable from the beginning. Twenty of the 24 identified long-term observable OTUs including two of the predictive OTUs, belong to the Firmicutes. Among them are OTUs of *Lactobacillus* that are thought to be associated with a healthy gut (31). In particular, *L. johnsonii* isolated from BALB/c mice has been shown to protect the host by influencing the host immune response against airborne allergens and viral infection (32). In our study, the proportional decrease in *L. johnsonii* (OTU0255) at ‘OD’ and ‘VO’ life stage is consistent with previous studies of aged mice (141 weeks old) and humans (76 to 100 years old) and may indicate a reduced protective effect from this species (18, 33). Among the few OTUs (e.g. *Corynebacterium* (OTU0822)) that emerged in the late phases of murine life (Supplementary Table S1), *Corynebacterium* spp. have been shown to also increase in relative abundance with age in humans from 79 years old and above (34). Hence, ‘OD’ and ‘VO’ C57BL/6J mice might be suitable models for studies related to infections from potentially pathogenic *Corynebacterium* spp. (35). *Faecalibaculum rodentium* Alo17, previously isolated from a 36-week old female C57BL/6J mouse, was shown here to represent the most abundant OTU in adult mice (Figure 3) (36). Altogether, these results suggest that the microbiome gradually changes throughout the lifespan of the animals and that OTUs of different phyla are affected to a varying degree, which in turn results in characteristic life-stage dependent microbiome configurations.

The random forest regression model identified key OTUs, including some low-abundance OTUs, that are predictive of host age (Figure 2c, Supplementary Figure S5 and Supplementary Table S2). Specifically, OTU0928 (*Parasutterella* sp. of Proteobacteria) was most predictive for the temporal changes with succession at ‘MD’ stage of the mouse life (Figure 2c). *Parasutterella* is ubiquitous in the gut microbiomes of mammalian and human hosts and may benefit the host with bile acid maintenance and cholesterol metabolism (37). The successional pattern observed for most Proteobacteria OTUs to be mid to late successors is consistent with other murine and human gut microbiome studies (2, 18, 27, 38). In contrast, *Desulfovibrio* sp. 16x (OTU0236), a sulfate reducing bacterium, represented an exception to the Proteobacteria successional pattern: This OTU was detected from ‘ MA’ to ‘ VO’ stages (Figure 2c), supposedly performing an important role along with acetogens in lowering hydrogen partial pressure in the gut (39, 40). Notably we did not detect any *Enterobacteriaceae*, however, this may be due to the commercial source of the laboratory mice in this study (41).

Our study clearly demonstrates the continuous changes of the gut microbiome and the different OTU succession patterns. The gut microbiomes of C57BL/6J mice largely followed host physiological development, but the periods of transition varied (10). For example, the transition from ‘MA’ to ‘MD’ was much longer (21 weeks) than the transition from ‘MD’ to ‘OD’ (5 weeks). The reliable host age prediction showed that there was sufficient distinction among few ‘source’ microbiota to delineate the majority of fecal microbiomes from ‘ MR’ to ‘ OD’. Limits to the prediction were observed for mice between the ‘OD’ and ‘VO’ stage, where the microbiota were too similar to be differentiated. The natural lifespan of mice sets limits to extend these experiments and to observe more small-scale dissimilarities between microbiomes of the very late stages. It also needs to be noted that these experiments were conducted in a well-controlled laboratory environment and the same standard laboratory feed was used for the entire duration. Using a different diet, alternating between diets, or performing other experimental modifications may also have lasting effects on the microbiome composition and its association with age as previously reported (18). Nevertheless, this study highlights the age-associated natural variation of the gut microbiome of conventionally raised C57BL/6J mice.

In summary, this study shows that the fecal microbiome of laboratory mice changes substantially throughout the entire adult age. Consequentially, this has implications for the design of experiments where the microbiome can be considered a contributing factor affecting host physiology. Furthermore, this study highlights that the microbiome can serve as a biomarker of aging and that host age can be inferred from microbiome composition.

## Materials and methods

### Animal husbandry, fecal sampling and DNA extraction

Experiments involving mice were approved by the Institutional Animal Care and Use Committee (IACUC number: TLL-17-018) in accordance with National Advisory Committee for Laboratory Animal Research guidelines and were performed at Temasek Life Sciences Laboratory, Singapore with supervision by trained veterinarians. Male C57BL/6J mice were purchased from InVivos (Singapore) at 63 days of age. The mice (*n* = 20) were kept in four cages of five mice each and maintained on standard chow (carbohydrate = 62.3%, protein = 25.5%, fat = 13.1%; PicoLab^®^ Rodent Diet 20; LabDiet, St Louis. MO, USA) *ad libitum*. Fecal materials were taken from all mice at one (10 weeks old) and two-week (12 weeks old) intervals followed by monthly (4-5 weeks old) intervals from 17 to 112 weeks old except upon arrival (9 weeks old) where a sub-sample of mice (*n* = 4) were sampled (Figure 1). Fecal matter was collected fresh in 2 mL sterile screw-capped tubes and flash-frozen in liquid nitrogen before storing at −80 °C until DNA extraction. A bead-beating phenol chloroform DNA extraction method was used on all fecal samples as previously described (42).

### Amplicon sequencing of 16S rRNA genes

A dual indexed 16S rRNA gene amplicon library was generated using primers 515F (43) and 806R (44) in triplicate PCRs per sample according to the protocol and indexes described in the Earth Microbiome Project (45). Illumina MiSeq sequencing was performed at the Genome Institute of Singapore according to the MiSeq reagent kit v2 (2 × 250 bp) preparation guide (Illumina, San Diego, CA, USA).

### Sequence processing and microbiota analysis

All MiSeq fastq files were deposited in the NCBI Sequence Read Archive under the Project ID PRJNA503299. The fastq files were processed using QIIME 2 version 2020.2 (accessed on 15 February 2020) using “qiime tools import” (46). Default options were used for all QIIME 2 scripts unless stated otherwise. Forward reads were quality denoised, trimmed, *de-novo* clustered and chimera checked using the “qiime dada2 denoise-single” command for DADA2 (47) with the following options: “--p-trim-left 23”, which removes the first 23 bp of reads and “--p-trunc-len 199”, which truncates at 199 bp. The quality filtered reads of 176 bp (*n* = 433) have a median count of 13,596 reads. To reduce the number of highly similar sequences, denoised single-end reads were clustered at 99% similarity OTUs using the “qiime vsearch cluster-features-de-novo” command with “--p-perc-identity 0.99” option, (48) which generated 1,107 OTUs. A single OTU for chloroplast was filtered prior to further analysis. Alignment was performed using the “qiime phylogeny align-to-tree-mafft-fasttree” command. The command “qiime diversity core-metrics-phylogenetic” with the option “--p-sampling-depth 3422” was used to generate alpha and beta diversity measures including Bray-Curtis dissimilarity, weighted- and unweighted-UniFrac matrices, PCoA plots and a rarefied OTU table, which were visualized as Emperor plots (49). Custom PCoA plots were generated using the “qiime emperor plot” command to plot samples grouped by age on the x-axis against the first principal coordinate (PC1) on the y-axis. Alphadiversity, relative abundance and heatmap plots were generated using the phyloseq (50), ggplot2 (51), reshape (52), microbiome (53), genefilter (54), data.table (55) and patchwork (56) packages for R (57). Taxonomic identities were assigned to OTUs using the “qiime feature-classifier classify-sklearn” command against a trained classifier SILVA SSU for V4 region version 132 non-redundant 99% identity database (58). To obtain updated taxonomic identities not curated in SILVA version 132, OTUs were annotated to the GenBank non-redundant database (accessed 28 Feb 2020) using the megablast function of BLASTn version 2.10.0+ (59, 60). To identify OTUs that are predictive of the temporal changes, the “qiime longitudinal feature-volatility” command was used with “--p-n-estimators 100” and “--p-random-state 10” options that adopts the random forest regressor as a machine learning method (14, 61). SourceTracker version 0.9.1 was used to estimate the mouse age using the rarefied table (15). As SourceTracker is estimating the probability of a ‘sink’ microbiota compared to a “source” microbiota, samples picked as ‘source’ were based on the timepoints with the fewest number of significantly different pairs i.e. more similar microbiome to most samples using the pair-wise permutational analysis of variance (PERMANOVA) analysis of the Bray-Curtis matrix (Supplementary Figure S6). One representative timepoint was picked as “source” for each of the phases with three or fewer timepoints namely, ‘MR’, ‘MA’ and ‘VO’. If all timepoints within each phase shared the same number of significantly different pairs, the median point was selected. Two timepoints were selected as “source” for the other two phases. All samples including those selected as “source” were also analyzed as “sinks”.

### Statistical analysis

The Kruskal-Wallis and pairwise Wilcoxon rank sum tests were performed using phyloseq (50), tidyr (62) and dplyr (63) packages for R (57). Pair-wise PERMANOVA tests were performed using the “qiime diversity beta-group-significance” QIIME 2 command based on 9,999 permutations with *P*-values corrected using the Benjamini-Hochberg FDR method (64). Spearman correlations and polynomial regression for SoureTracker prediction between actual and predicted ages were performed using R (57). *P*-value < 0.05 is considered statistically significant.

## Acknowledgements

We thank and Muhammad Khairillah Bin Nanwi at TLL biocomputing for bioinformatics support. We also thank Subramanian Kabilan and Adeline Wong for support with animal maintenance and sample collection, respectively.

---

**Supplementary Figure S1.** Survival plot of male C57BL/6J mice throughout the study. The red line indicates when the mice arrived at the housing facility and when the first fecal samples were collected. Longer tick marks on x-axis indicate sampling time points.

**Supplementary Figure S2.** Longitudinal changes in alpha-diversity analysis of fecal microbiota of 9-to 112-week-old C57BL/6J mice (*n* = 4-20). All alpha-diversity measures are based on an OTU table rarefied to 3,422 counts per sample. ‘MR’ (weeks 9-12; *n* =4-19), ‘MA’ (weeks 17-22; *n* =19), ‘MD’ (weeks 43-60; *n* =15-20), ‘OD’ (weeks 82-104; *n* =15-18) and ‘VO’ (weeks 108-112; *n* =12-13).

**Supplementary Figure S3.** Principal coordinate analysis (PCoA) plots based on beta-diversity measures. PCoA plots showing PC1 relative to age of **(c)** Bray-Curtis dissimilarity and **(d)** weighted-UniFrac distance matrices. Percentage variation is shown in parenthesis. Beta-diversity measures were based on an OTU table rarefied to 3,422 counts per sample. OTUs were clustered at 99% similarity cut-off.

**Supplementary Figure S4.** Longitudinal changes in mean relative abundance of less abundant phyla (mean relative abundance ≤3%). Relative abundances are based on an OTU table rarefied to 3,422 counts per sample.

**Supplementary Figure S5.** Normalized counts of predictive 28 OTUs not among the top 110 OTUs using a random forest regression method. Normalized counts are based on an OTU table rarefied to 3,422 counts per sample. Taxonomic assignments >99% nucleotide identity for species and 95%-99% identity for genus level were based on top BLASTn hits and <95% nucleotide identity for family level were based on the SILVA SSU database 132 release.

**Supplementary Figure S6.** Beta-diversity measures and pairwise PERMANOVA test between timepoints. The weighted-UniFrac distances of five timepoints of each life phase relative to the other time-points **(a)**. The five time-points arranged from top to bottom are 12 weeks old, 22 weeks old, 56 weeks old, 91 weeks old and 108 weeks old. Red dot and red line denote FDR-corrected *P*-values based on pairwise PERMANOVA test using 9,999 random permutations. An FDR-corrected *P* < 0.05 is considered statistically significantly different. Horizontal dotted line indicates the FDR-corrected *P* = 0.05 cut-off. A summary of total count of significantly different pairs of microbiota across the study period relative to each timepoint for Bray-Curtis dissimilarity and weighted UniFrac matrices **(b)**. Dotted line indicates the line of best fit over the 26 timepoints for each beta-diversity matrix. Black dots on Bray-Curtis plot denotes the points used as ‘source’ for SourceTracker.

